# In transected nerves, distal repair Schwann cells are required at the injury site to direct and accelerate axonal regrowth

**DOI:** 10.64898/2026.02.17.706483

**Authors:** Daniel E. Lysko, Annika R. Johnson, William S. Talbot

## Abstract

In vertebrate peripheral nerves, damaged axons can regrow after injury, but the outcomes of regeneration are variable and often incomplete. Schwann cells in injured nerves are important for repair, but their actions at different positions and stages of nerve repair are not well understood. We have investigated the roles of Schwann cells in a larval zebrafish nerve injury model, in which nerves are visible in living animals during development, the initial injury response, and regrowth of the transected axons. After mechanical injury, distal Schwann cells adopt a repair phenotype characterized by changes in marker expression, elongation, and ability to guide axons across the injury site. In contrast, proximal Schwann cells are not sufficient to guide axons across the injury site, and they associate with axons that regrow along aberrant paths. In *erbb2* mutants lacking Schwann cells, developmental axon growth is normal, but after transection, axonal regrowth is greatly slowed and often misdirected. By examining animals with nerves partially populated by Schwann cells, we find that axons can regrow through regions devoid of Schwann cells, provided that at least one distal Schwann cell is at the injury site. Timelapse imaging reveals that distal Schwann cells extend processes toward the injury site, which contact and guide axons regrowing from the proximal nerve stump. In *irf8* mutants lacking macrophages, debris from transected axons is cleared on schedule, and axonal regrowth is normal. Our studies demonstrate that Schwann cells immediately distal to the injury site have a unique and essential role in axonal regrowth.

**Main Points:**

- After nerve transection in larval zebrafish, proximal and distal Schwann cells have distinct functions at injury site
- A single distal repair Schwann cell is sufficient for axonal regrowth
- Axonal regrowth is normal in mutants without macrophages

## Introduction

Elucidating the mechanisms that control the repair of the nervous system after injury remains a great challenge (Jessen and Mirsky, 2016). One challenge is the nature of injury, which may vary widely in region, scope, and life stage. For example, repair of a nerve in an adult may involve regrowth of axons over distances much greater than those in the embryo, and at stages when the developmental guidance cues are no longer operative. Furthermore, the debris and inflammation generated by injury may impede regrowth and repair (Rotshenker, 2011). On the other hand, injury occurs in the context of a pre-existing, functional nervous system, which may provide a template for repair.

Schwann cells in peripheral nerves adapt to injury of their associated axons and promote axonal regrowth, providing a compelling example of cellular redeployment after injury (Arthur-Farraj et al., 2012). After a nerve is transected, the distal regions of severed axons degenerate (Waller, 1851; Beirowski et al., 2005). The denervated Schwann cells greatly alter their functions to adopt a distinct cell state that promotes repair (Arthur-Farraj et al., 2012, 2017). During development and homeostasis, Schwann cells make and maintain myelin, but in stark contrast repair cells degrade myelin and engulf axonal debris, thereby clearing the distal nerve segment to allow axonal regrowth (Jessen and Mirsky, 2016; Jessen and Arthur-Farraj, 2019). Repair cells also attract macrophages, which may contribute to debris clearance after injury (Perry et al., 1995; Dailey et al., 1998; Villegas et al., 2012; Morales and Allende, 2019). In mammals, repair Schwann cells greatly increase their length, forming characteristic structures called Büngner bands that provide a substrate for axons regrowing through the distal nerve segment (Gomez-Sanchez et al., 2017). Thus, injury reverses the relationship between Schwann cells and axons: axons guide Schwann cell migration during development but are apparently guided by repair Schwann cells during regrowth (Gilmour et al., 2002; Ceci et al., 2014; Jessen and Mirsky, 2016).

Many studies demonstrate that Schwann cells are essential for nerve regeneration after injury (Jessen and Mirsky, 2016; Bosch-Queralt et al., 2023). In zebrafish mutants lacking Schwann cells, motor and sensory axons are relatively normal during initial outgrowth, but axonal regrowth after injury is delayed, misrouted, and incomplete (Gilmour et al., 2002; Lyons et al., 2005; Perlin et al., 2011; Villegas et al., 2012; Ceci et al., 2014; Rosenberg et al., 2014; Xiao et al., 2015; Walker et al., 2023). In mammals, c-Jun is essential for Schwann cells to convert to the repair state, and conditional c-Jun mutants have diminished myelin clearance, disrupted axonal regrowth, and impaired functional recovery (Arthur-Farraj et al., 2012; Ruff et al., 2012). Although repair Schwann cells play key roles following injury, important questions remain about the distinct functions of Schwann cells in different positions along the injured nerve, including the distal and proximal nerve segments and the injury site itself (Graciarena et al., 2014; Xiao et al., 2015; Clements et al., 2017). Furthermore, the requirements for macrophages that are recruited to injured nerves remain to be clarified.

In the present study we have developed a mechanical nerve injury model in the larval zebrafish, which is genetically tractable and allows in vivo imaging of nerves before, during, and after nerve injury and repair. Complementing prior models using laser and electroablation (Rosenberg et al., 2012; Villegas et al., 2012; Moya-Díaz et al., 2014; Tian et al., 2023), our mechanical injury model is rapid, reproducible, and accessible. We find that Schwann cells distal to the injury site display characteristics of repair Schwann cells previously defined in mammals. Repair Schwan cells elongate and extend processes that contact and guide axons regrowing across the injury site (Stoll and Müller, 1999; Gomez-Sanchez et al., 2017; Walker et al., 2023). We found that axonal regrowth is impaired in the absence of any distal Schwann cells, but that extensive regrowth can occur when only a single distal Schwann cell is present. Finally, we found that axonal regrowth was normal in mutants lacking macrophages. Our studies demonstrate that distal repair Schwann cells at the injury site have unique and essential functions in axonal regrowth.

## Methods

### Zebrafish lines and maintenance

All zebrafish experiments were conducted using protocols approved by the Stanford University institutional animal care and use committee and conforming to appropriate city, state, and national regulations. Embryos and larvae were anesthetized with 0.016% Tricaine (Syncaine, Syndel) prior to experimental procedures. Embryos and larvae were treated with 1-phenyl 2-thiourea (PTU, 0.003%, Sigma) to inhibit pigmentation for microscopy. Embryos and larvae were analyzed up to 6 dpf. The *erbb2*^st61^ and *irf8*^st96^ mutant lines have been previously described (Lyons et al., 2005; Shiau et al., 2015).

### Nerve transection

Larvae were anaesthetized with Tricaine and mounted on their side on a coverslip in 0.9% agarose (UltraPure LMP Agarose, Invitrogen). The left posterior lateral line nerve was transected at approximately the seventh somite (+/- 1 somite) with an insect pin (stainless steel Minutien pins, 0.02mm tip, 0.2mm rod, FST). Immediately after injury the larvae were imaged to confirm full transection of the nerve using a Zeiss LSM700 confocal microscope. Nerves were visualized using the Tg(NBT:dsRed) line (Peri and Nüsslein-Volhard, 2008). During this initial injury screening, we excluded any larvae that did not have fully transected nerves and those that had excessive injuries, such as injury to the spinal cord or yolk. Schwann cells were visualized using the Tg(zFoxd3:GFP(17)) line (Gilmour et al., 2002). Remyelination was visualized with a Tg(mbp:GFP-CAAX) line (Almeida et al., 2011). Macrophages were visualized using the Tg(mpeg:GFP) line (Ellett et al., 2011).

### Generation of nerves partially covered with Schwann cells

Embryos were treated beginning at 22-23 hpf with 0.6 µM AG1478 (Cayman Chemical) in 1% DMSO (Chem-Impex). Embryo water containing this ErbB inhibitor was changed daily. In different experiments, treatment with the inhibitor was either continued throughout injury and recovery, or removed at the time of injury to allow nerve recovery in the absence of ErbB inhibitor. At 3 dpf, animals were mounted as above, the leading Schwann cell position was observed using epifluorescence microscopy, and the nerve was injured as above while choosing injury position to generate animals with either zero, one, 2-3 or many Schwann cell distal to the injury site. Due to variability in Schwann cell migration or effect of ErbB inhibition, the leading Schwann cell position varied (Figure 5B). Animals with leading Schwann cells positioned at somite 10-16 were selected, allowing 14-20 somites of possible regrowth. After injury, full nerve transection and the number of Schwann cells distal to the injury site were assessed using confocal microscopy. Animals imaged on multiple days were individually housed in 6-well plates.

### Axonal regrowth rate calculation

NBT:dsRed transgenic larvae were mounted, imaged, unmounted and imaged again at successive stages during development and axonal regeneration. Developmental growth rate was determined by measuring the entire length of the nerve at two different time points in the same individual and dividing that length by the number of hours between imaging time points. Axonal regrowth rate was similarly determined; in some cases the length of the nerve from the most proximal part of the nerve which was unaffected by the injury was used as the landmark from which to assess the length of total regrowth. In the case of *erbb2* mutants, in which the site of injury was not easily determined at later stages because of axonal retraction and disrupted regrowth, the entire nerve was imaged so that the difference in length at the respective time points could be calculated. The rate in these cases was the difference in nerve length divided by the number of hours between the start of the two imaging sessions. Image montages were assembled using the plugin MosaicJ (Thévenaz and Unser, 2007) in Fiji/ImageJ (Schindelin et al., 2012).

### Rate of axonal regeneration before and after contacting a distal Schwann cell process

To quantify the effects of distal Schwann cell contact on axonal regeneration, we analyzed one time lapse movie in which the time of initial contact was evident (Supplemental Movie 1). Using the ImageJ line segment tool, we marked a spot slightly behind the growth cone in each frame and advanced frame by frame for the desired time window. Once the path length of regrowth was known, it was divided by the time interval. When the nerve was obstructed or out of frame at the precise time point, we adjust the time to a point when the growth cone was again visible. Because the individual axons were sometimes thin and dim, it was necessary to greatly increase the saturation to visualize the most distal axon in the nerve at each timepoint.

### Quantification and statistical analysis

Graphs and statistical analysis were performed in GraphPad Prism. Non-normal distributions were compared using the Mann-Whitney test. Exact p values are reported.

## Results

### Mechanical nerve injury model in larval zebrafish

Building on prior studies that employed electroablation and laser ablation of the larval zebrafish posterior lateral line nerve (PLLN), we sought to develop a mechanical transection model that was fast and reproducible with high recovery and survival rates. To observe neurons and Schwann cells, we used the transgenes NBT:dsRed and foxd3:GFP, respectively (Fig. 1A TG) (Gilmour et al., 2002; Peri and Nüsslein-Volhard, 2008). During the initial phase of posterior lateral line development, the lateral line primordium migrates in the skin along a roughly linear path from the ganglion to the tip of the tail, starting at about 22 hours post fertilization (hpf) and concluding at about 40 hpf (Figure 1 B, C)(Kimmel et al., 1995; Nechiporuk and Knaut, 2025). The migrating primordium precedes and directs the outgrowing PLLN axons (Metcalfe, 1985; Gilmour et al., 2004; Grant et al., 2005), which in turn direct the migration of accompanying Schwann cells (Gilmour et al., 2002; Lyons et al., 2005). Schwann cells are required to shift the location of the axons from a superficial position in the skin to a deeper position beneath the skin basement membrane (Raphael et al., 2010). By 3 dpf, active neuromasts are innervated, and several myelinated axons are evident in the PLLN (Sarrazin et al., 2010; Raphael et al., 2011; Nechiporuk and Knaut, 2025).

**Figure 1.**
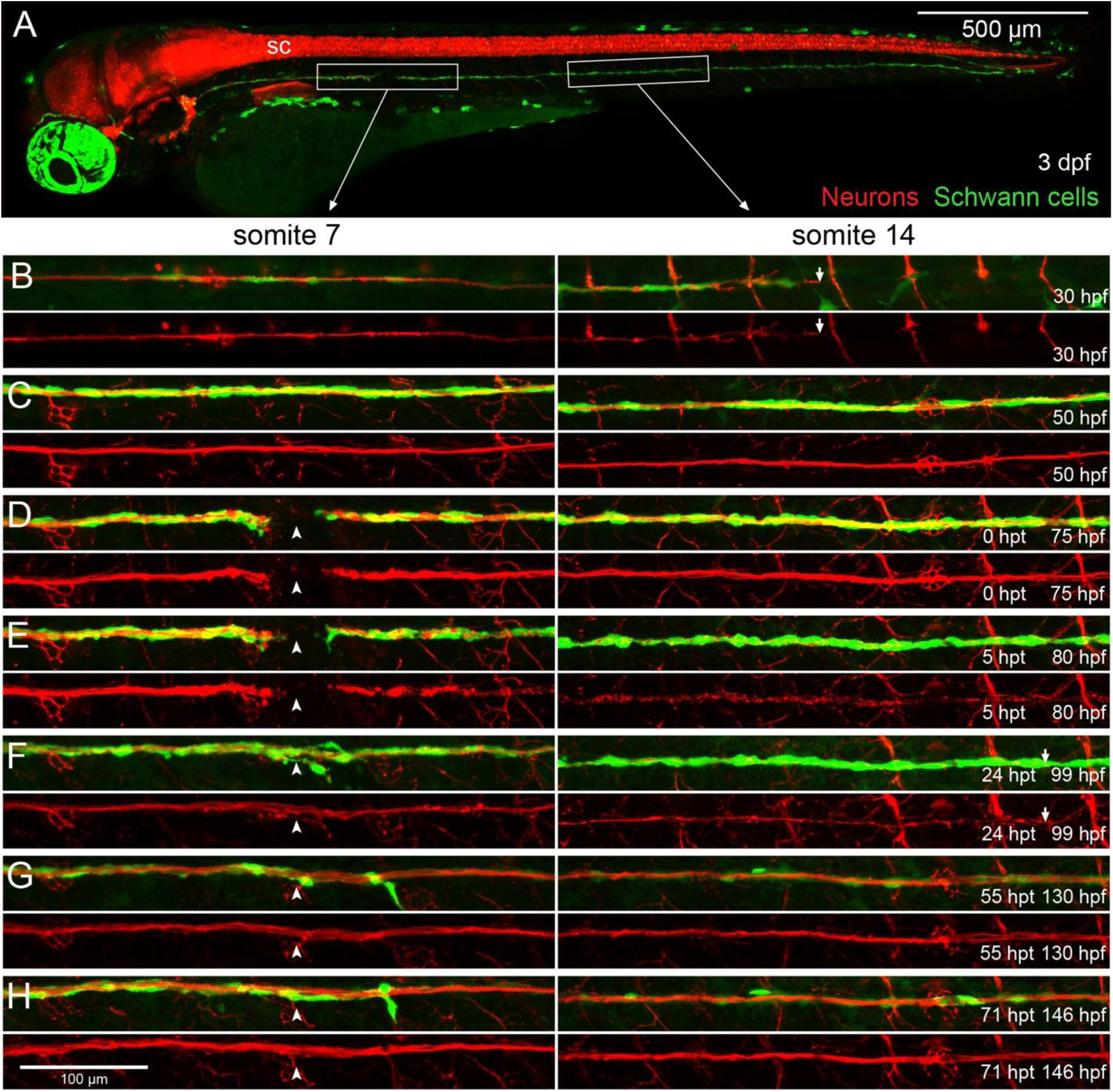
Time course of posterior lateral line development and recovery after transection in wildtype larvae. **(A)** Wildtype larva with transgenic markers of Schwann cells (foxd3:GFP, green) and neurons (NBT:dsRed, red) a few hours after mechanical injury of the PLLN at the region of somite 7 at 3 dpf. White boxes denote regions shown at higher magnification in subsequent panels. The spinal cord (sc) and other deep structures appeared intact. **(B-H)** Time course of images showing development and regrowth of PLLN after mechanical injury (arrowheads mark injury site). **(C-H)** Images of the same fish at the indicated stages. Different animals are shown in **(A)** and **(B)**. The tips of growing axons (arrows) are noted during development (**B**, 30 hpf) and regrowth (F, 24 hpt).

For our mechanical injury studies, we employed an insect pin to transect the PLLN at 3 dpf (Figure 1A, D). A small incision was made at the location of the horizontal myoseptum of somite 7 of anesthetized larvae mounted in agarose. The injured area included superficial muscle and skin cells in addition to the nerve itself. Soon after injury, confocal imaging was used to verify that the nerve was completely transected and that the spinal cord and other deep structures appeared to be intact. The gap in the transected nerve and the overall injured area typically spanned 50-100 micrometers.

As in prior studies (Villegas et al., 2012; Ceci et al., 2014), the distal nerve segment was fragmented and being cleared by 5 hours post transection (Figure 1E). Around this same time, axons began to project from the proximal nerve segment through the injury site. We monitored axonal regrowth either by reference to the somite position or by length of axon from the injury site. The distal nerve segment through which axons regrew was a path of approximately 23 somites (1850 µm) at 3 dpf. Almost all wildtype animals showed significant axonal regrowth along the original path of the nerve by 24 hours post-transection (hpt) (Figure 1F) and complete regrowth by 48 hpt (Figure 1 G, H). As noted in prior studies (Ceci et al., 2014), Schwann cells remained in both the proximal and distal nerve segments throughout regrowth, and the expression of the foxd3:GFP transgene was diminished specifically in the Schwann cells of the distal segment by 2 days post transection (dpt) (Figure 1G, H). This differential expression prompted us to further investigate other differences between proximal and distal Schwann cells.

### Distal Schwann cells exhibit characteristics of repair Schwann cells

Although the axons in almost all wildtype animals had completely regrown by 48 hpt (Figure 1), there was variability at the injury site, as evidenced both by the variable time proximal axons took to cross to the distal segment and the number of misrouted proximal axonal branches crossing the injury site in different animals (Figure 2A-C). Interestingly, the misrouted axonal branches associated with proximal Schwann cells (evident from strong expression of foxd3:GFP), whereas the axons regrowing along the proper path associated with the distal Schwann cells with diminished foxd3:GFP (Figure 2A-C). Thus, these results suggested that the distal Schwann cells can guide regrowing axons along the proper path—a function reminiscent of repair Schwann cells in mammals (Thomas and King, 1974; Son and Thompson, 1995; Arthur-Farraj et al., 2012; Cattin et al., 2015). As observed for mammalian repair Schwann cells (Arthur-Farraj et al., 2012, 2017; Brosius Lutz et al., 2022), distal Schwann cells down-regulate the myelination reporter *myelin basic protein:*GFP (*mbp:*GFP); expression was diminished at 1 dpt and 2 dpt, but was returning by 3 dpt (Fig. 2D). After electroablation of the PLLN, Ceci et al. (2014) showed that expression of Mbp protein was reduced at 48 h post injury. Although both the mbp:GFP and foxd3:GFP transgenes demonstrate transcriptional changes in presumptive repair Schwann cells in the distal segment of the PLLN, these reporters follow different time courses. The down regulation of foxd3:GFP was only apparent at 2 dpt, and it continued to decrease at 3 dpt, when mbp:GFP was reactivated as the regrown axons become remyelinated by distal Schwann cells.

**Figure 2.**
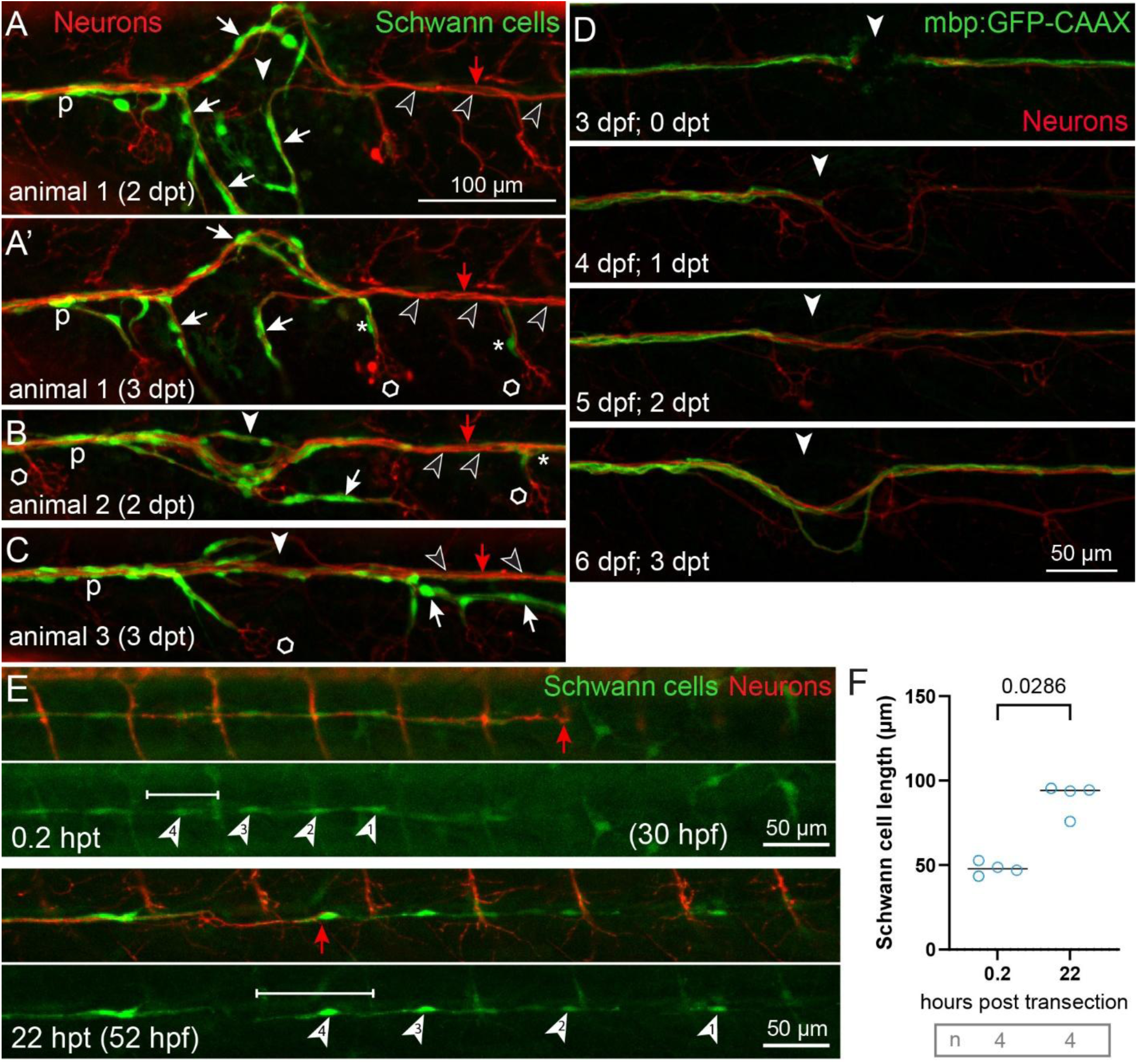
Distal Schwann cells display characteristics of repair Schwann cells. (A-C) Three different examples of axons regrowing at the site of mechanical injury (white arrowhead). The anterior (proximal) side of the nerve is to the left in all images. Although all Schwann cells express the foxd3:GFP transgene (green), proximal Schwann cells (p) continue to highly express foxd3:GFP, while by 2 dpt, distal Schwann cells (black arrowheads) have weak expression. Axons regrowing along the correct path (red arrow) associated with distal Schwann cells that weakly express the reporter, whereas proximal Schwann cells associated with misrouted axons (white arrows). (A’, B) By 2-3 dpt, some medium expressing Schwann cells(*) associated with axons projecting from the main nerve trunk and innervating neuromasts (hexagons). Neurons are labeled with the NBT:dsRed transgene (red). **(D)** Time course of expression of mbp:GFP-CAAX myelin reporter (green) and neuronal reporter NBT:dsRed (red) after mechanical injury and nerve regrowth. By 1 dpt, axons have regrown through the injury site (arrowhead), and myelin reporter expression was greatly reduced in the distal segment (right side of the image). By 3 dpt, myelin reporter expression had recovered. **(E)** Growing PLLN axons and Schwann cells at 30 hpf, 0.2 hours after transection. The injury site is anterior (proximal) to the region shown. Arrowheads note four different Schwann cells near the growing tip of the nerve. These same cells are shown 22 hours post transection (hpt), when axons are regrowing into the region (arrow) and the Schwann cells have greatly increased their length (for example cell 4; brackets), as quantified in **(F)**.

Mammalian repair Schwann cells are much longer than immature and myelinating Schwann cells (Gomez-Sanchez et al., 2017), so we sought to measure the length of presumptive repair Schwann cells in the distal PLLN after injury. After PLLN outgrowth is complete, the number and density of Schwann cells in the nerve impede visualization of individual cells using foxd3:GFP (Figure 1) (Gilmour et al., 2002; Ceci et al., 2014). We found that individual Schwann cells can be visualized after injury at an earlier timepoint (30 hpf), so we measured the length of Schwann cells migrating near the tip of the growing PLLN (Figure 2E), soon after nerve transection and 22 hours later (Figure 2E and 2E’). By this later timepoint, distal axons had degenerated and axonal regrowth had begun (Figure 2E’). By 22 hpt, distal Schwann cells had lengthened significantly (Figure 2E’, 2F). These results indicate that at 1-2 dpt distal Schwann cells greatly elongate, downregulate myelin genes, and guide regrowing axons—all key features of repair Schwann cells as described in mammals. Furthermore, our results suggested that there are functional differences in the ability of proximal and distal Schwann cells to guide regrowing axons.

### Schwann cells are essential for normal axonal regrowth after transection

Previous analyses of laser ablation and electroablation of motor nerves and PLLN have demonstrated that Schwann cells have a key role in guiding regrowing axons, but their role in stimulating axonal regrowth is less clear (Villegas et al., 2012; Ceci et al., 2014; Rosenberg et al., 2014; Xiao et al., 2015). To address these questions in our mechanical injury model, we analyzed *erbb2* mutants, which are devoid of Schwann cells (Figure 3). As in previous studies (Gilmour et al., 2002; Lyons et al., 2005), developmental outgrowth of PLLN axons appeared normal in *erbb2* mutants despite the lack of Schwann cells (Figure 3 B, C; Figure 4). In contrast to wild-type, in *erbb2* mutants both the proximal and distal segments of the injured nerve rapidly retracted away from the site of injury (Figure 3D). Axonal regrowth was greatly delayed in *erbb2* mutants (Figure 3 F, G; Figure 4). Even by 2 dpt, when axons had fully regrown in almost all wild-type animals (∼23 somites), axons in *erbb2* mutants typically had regrown only 1 or 2 somites past the injury site, and the rate of regrowth was approximately one-sixth of wild-type (Figure 4). Although Schwann cells can engulf degenerating axons (Catenaccio et al., 2017; Vaquié et al., 2019), we found that axonal debris was mostly cleared by 5 hpt in *erbb2* mutants, a timeline comparable with wild-type clearance (Figure 3E). These results indicate that Schwann cells are essential for regrowth of axons after mechanical injury.

**Figure 3.**
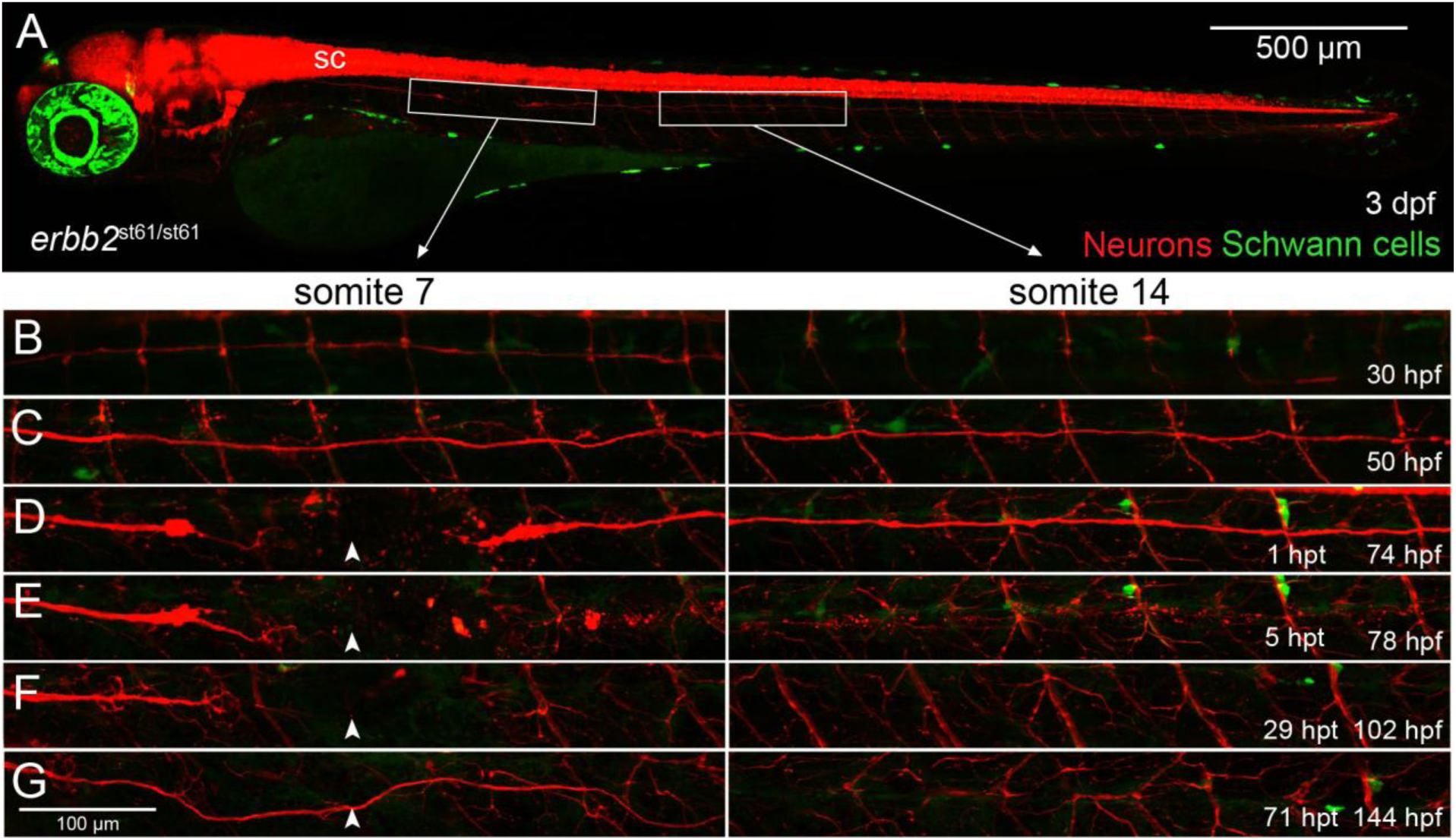
Time course of posterior lateral line development and recovery after transection in *erbb2* mutant. **(A)** Homozygous *erbb2* mutant larva with transgenic markers of Schwann cells (foxd3:GFP, green) and neurons (NBT:dsRed, red) one hour after mechanical injury of the PLLN at the region of somite 7 at 3 dpf. White boxes denote regions shown at higher magnification in subsequent panels. The spinal cord (sc) and other deep structures appeared intact. **(B-G)** Time course of images showing development and regrowth of PLLN after mechanical injury (arrowheads mark injury site). All images are from the same animal at the indicated stages. No Schwann cells were evident along the PLLN, and very few cells expressed the foxd3:GFP reporter in the mutant. The genotype was determined by PCR after imaging.

**Figure 4.**
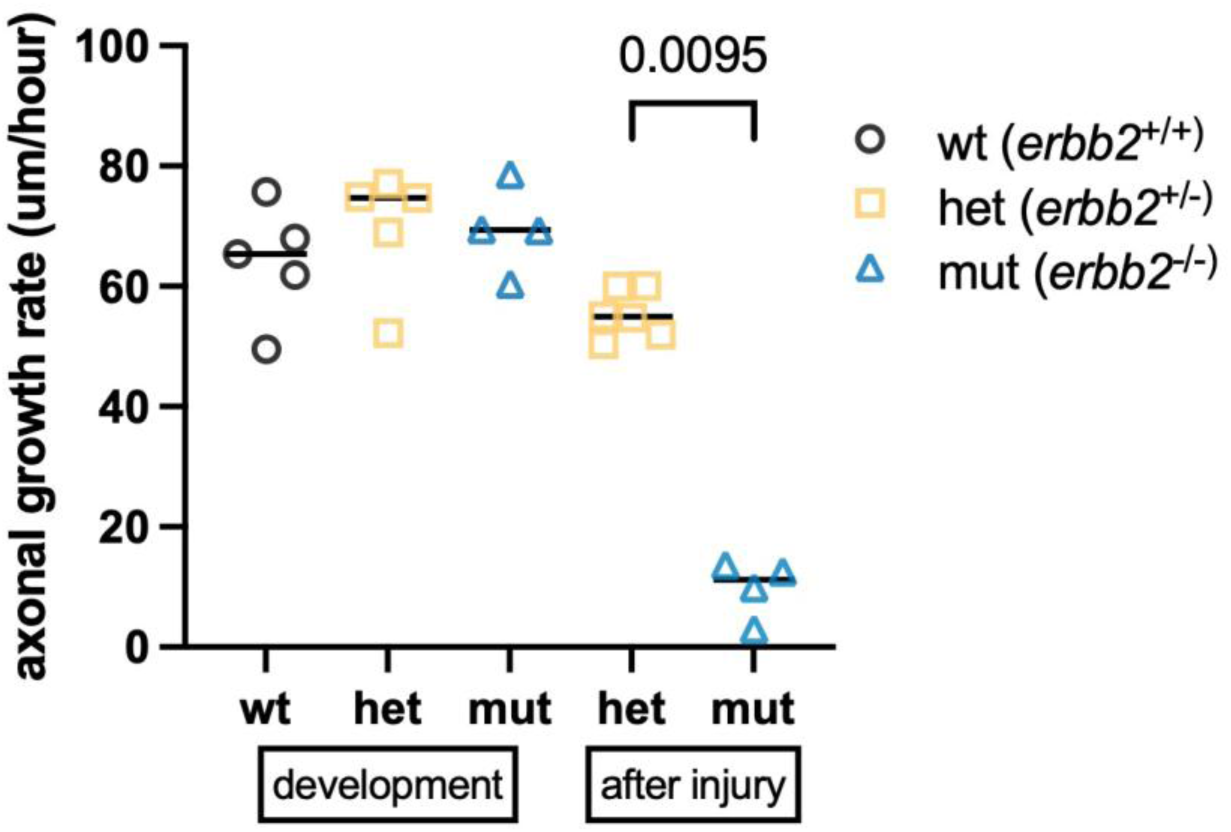
Rate of axonal outgrowth during development and regrowth after transection in wildtype and *erbb2* mutant larvae. Quantification of axonal growth rate in *erbb2* homozygotes compared to wildtype and heterozygous siblings. Rate of developmental outgrowth was calculated from 26-32 hpf, and the rate of regrowth was calculated from 1-2 dpt, after injury on 3 dpf. At the appropriate stages, axon lengths were measured in composite images of the PLLN. The genotypes of all animals were determined by PCR after imaging. Each point represents one nerve in one fish, measured at two timepoints to calculate the change in length over the intervening time. Lines indicate the median. Mann-Whitney test used for comparisons.

### A single Schwann cell on the distal side of the injury site is sufficient for extensive axonal regrowth

Distal to the injury site, repair Schwann cells may support axonal regrowth by providing both stimulatory signals and a permissive substrate (Son and Thompson, 1995; Yan et al., 1995; Parrinello et al., 2010; Rosenberg et al., 2014; Isaacman-Beck et al., 2015). These repair Schwann cells could mediate axonal regrowth mainly by facilitating regrowth across the injury site, or they may function along the entire distal nerve segment. To determine where repair cells function during axonal regrowth, we used ErbB inhibitors to generate animals with PLLN axons only partly covered with Schwann cells. Pharmacological inhibition of ErbB signaling decouples axonal growth from Schwann cell migration (Lyons et al., 2005), and addition of ErbB inhibitor after nerve outgrowth has commenced results in nerves partially populated with Schwann cells (Figure 5A). We measured the position of the most distal Schwann cell in individual ErbB-inhibited larvae (Figure 5B), and chose the injury site so that zero, one, a few, or many Schwann cells were located on the distal side of the transected nerve.

**Figure 5.**
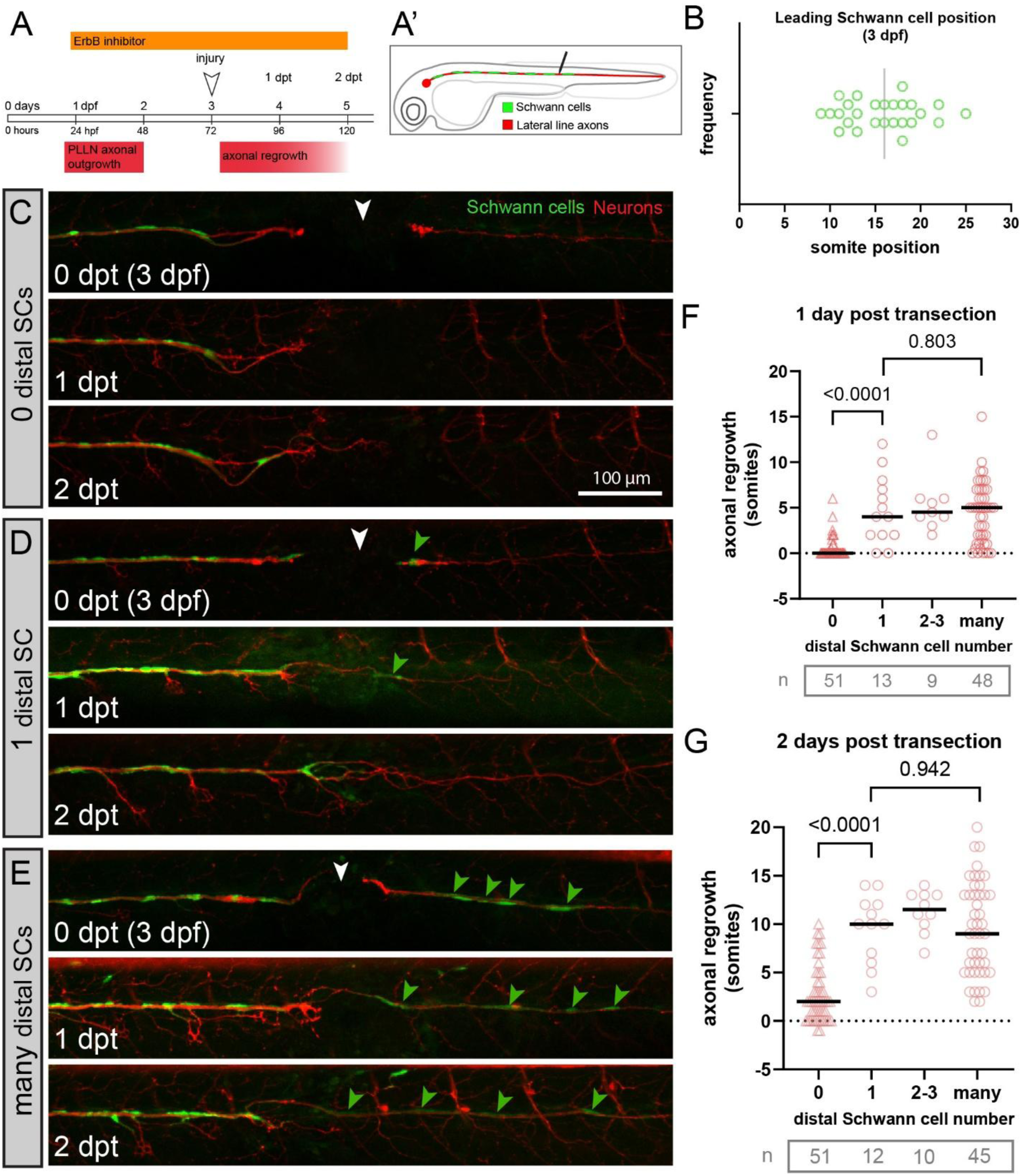
A single distal Schwann cell is sufficient for significant axonal regrowth after nerve transection. **(A)** Timeline of ErbB inhibitor treatment to generate nerves partially populated with Schwann cells. **(A’)** Transection position chosen to generate an injury site with a single distal Schwann cell. **(B)** Treatment with 0.6 micromolar AG1478 beginning at 23 hpf disrupts Schwann cell migration and results in a variable extent of Schwann cell position along the PLLN at 3 dpf. (C-E) Animals expressing foxd3:GFP (Schwann cells, green) and NBT:dsRed (neurons/axons, red) were longitudinally imaged from injury through 2 days post transection (white arrowhead at injury site, green arrowheads mark distal Schwann cells). **(C)** When the injury site was positioned to leave no distal Schwann cells, axons stall at the injury site 1 dpt. Only modest growth was observed at 2 dpt. **(D)** With a single distal SC, significant axonal growth was apparent at 1 dpt, and growth continued at 2 dpf. **(E)** When many distal Schwann cells were present, the extent of axonal regrowth was similar to **(D)**. **(F,G)** Quantification of axonal regrowth at 1 dpt and 2 dpt, when 0, 1, 2-3, or many distal Schwann cells were present. Lines indicate the median. Data from four independent experiments, n=animals. Mann-Whitney test used for comparisons.

In animals with no distal Schwann cells, transected axons typically stalled at the injury site at 1 dpt; and they made little progress by 2 dpt (Figure 5C, F, G); these results are very similar to *erbb2* mutants (Figure 3). Strikingly, even a single distal Schwann cell was sufficient to promote extensive axonal regrowth at 1 dpt and 2 dpt (Figure 5D, F, G). Animals with a few or many Schwann cells also showed extensive axonal regrowth, but the regrowth was not significantly better than animals with a single distal Schwann cell (Figure 5E, F, G). These experiments indicate that repair Schwann cells have an especially important function immediately distal to the injury site. These analyses also indicate that transected axons that are able navigate the injury site can subsequently regrow though large regions that are devoid of Schwann cells.

### Contact with the distal Schwann cell directs and accelerates axonal regrowth

We sought to further investigate the interactions between regrowing axons distal Schwann cells at the injury site using time-lapse imaging (Figure 6). The figure depicts frames from a time-lapse move that shows axons regrowing dorsally and ventrally from the injury site, beginning at the time when a distal Schwann cell process first contacts a ventral axon (Figure 6C, inset). Fifty minutes after this initial contact, a new axonal branch appeared at precisely the contact point (Supplemental movie 1). This new branch then followed the path formed by the distal Schwann cell process across the injury site and into the distal nerve segment, where it regrew along a path of Schwann cells (Figure 6D). Though this branch was formed later than many other branches that did not contact distal Schwann cells, it was the first to cross the injury site. After Schwann cell contact, the crossing axon appeared to accelerate as it regrew distally through the imaged region (Figure 6B, D). During the first 2 hours after Schwann cell contact, the axon grew at a rate of 30 micrometers per hour. By 10 hours after contact, the axon was regrowing at a rate similar to developmental outgrowth (compare Figure 6D to Figure 4), and more than double the growth rate of a misrouted axon that projected dorsally and was never contacted by a Schwann cell (Figure 6B). This analysis provides evidence that distal Schwann cell contact is important to direct regrowing axons across the injury site and accelerate their regrowth along the correct path.

**Figure 6.**
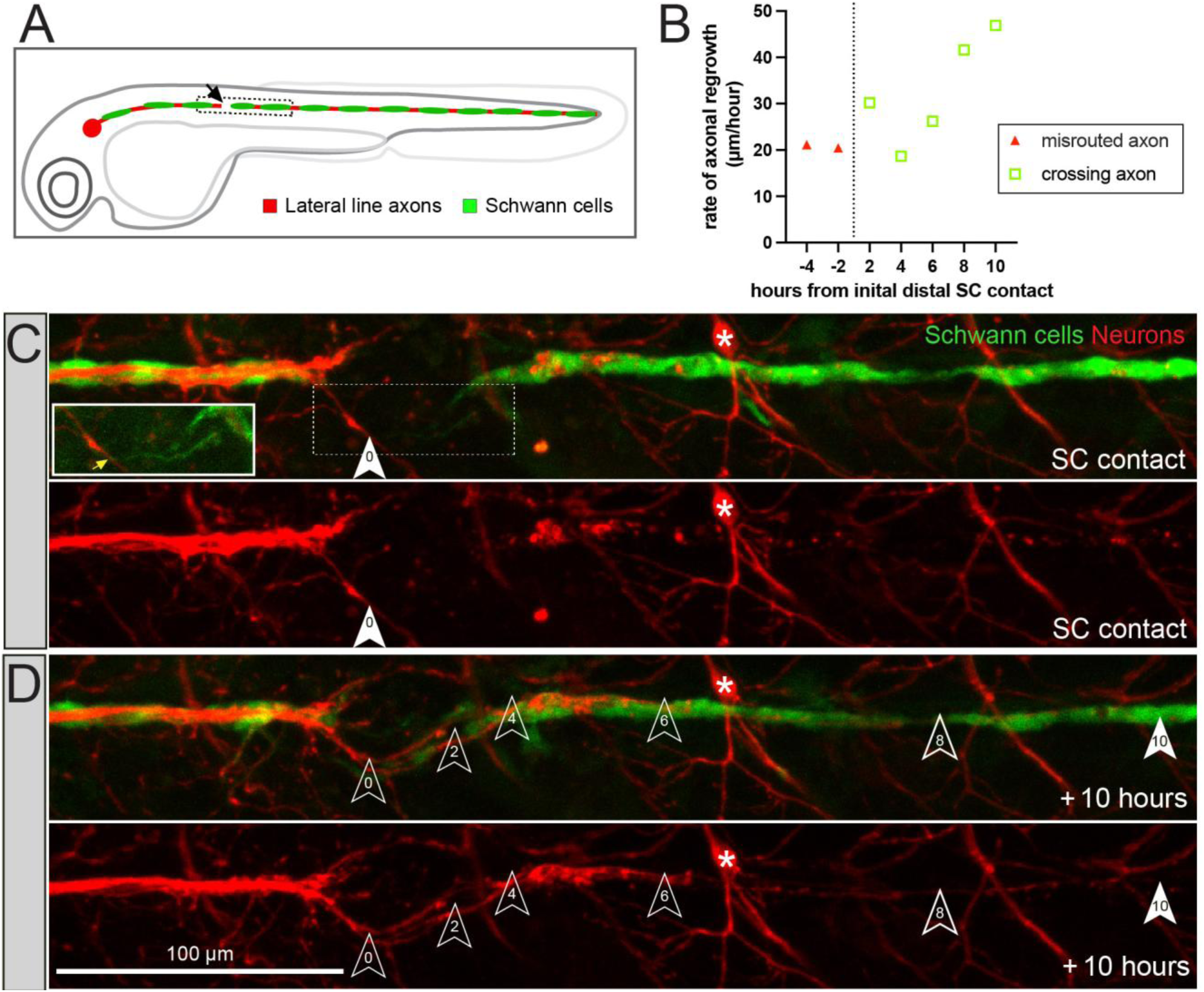
Time-lapse analysis of interaction between distal Schwann cells and axons regrowing after transection. (**A)** Diagram of larval fish with injury and imaging sites labeled. A wildtype larva with the Schwann cell and neuronal reporters was injured at 3 dpf, and analyzed with time-lapse imaging from 8 to 34 hpt. The initial point at which a distal Schwann cell process contacted a regrowing axon was determined, allowing analysis of axons before and after contact. **(B)** Quantification of growth rate of an early, mistargeted axon bundle which did not contact a distal Schwann and grew on an aberrant path (red triangles) and the axon bundle that was first contacted by a distal Schwann cell (green squares). The rate of axonal regrowth accelerated at 8 hours after Schwann cell contact. **(C)** Frames from the timelapse movie at the point when a distal Schwann cell first contacted a regrowing axon (arrowhead marked 0). Inset depicts brightened region denoted by dotted box, and yellow arrow marks Schwann cell-axon contact point). **(D)** Frame from the movie 10 hours after contact, showing that a new axon branch grew from the point of Schwann cell contact across the injury site and down the path of the PLLN. Arrowheads indicate the position of the most distal axon at 2-hour intervals after Schwann cell contact as indicated by the numbers in each arrowhead. Asterisk marks a motor neuron that is evident throughout the analysis.

### Normal axonal regrowth in the absence of macrophages

Schwann cells recruit macrophages to the injured nerve where they secrete cytokines and help phagocytose myelin and axonal debris (Perry and Brown, 1992). In addition to debris clearance, there is evidence that macrophage-endothelial cell communication can establish a cell bridge that guides regrowing axons of transected nerves (Cattin et al., 2015; Li et al., 2022). Nonetheless the requirement for macrophages for nerve growth is unclear (Villegas et al., 2012). To examine the role of macrophages directly, we employed *irf8* mutants, which lack macrophages (Shiau et al., 2015), while visualizing the macrophage response with the mpeg:GFP transgene (expressed in the myeloid lineage) (Ellett et al., 2011).

Axonal development in *irf8* mutants is indistinguishable from wild type (Figure 7A and B). In wild-type animals, a large number of macrophages occupied the injury site by 6 hpt (Figure 7C); most of these were no longer present by 2 dpt, when the transected axons had fully regrown (Figure 7G). In *irf8* mutants, only a few cells weakly expressing the mpeg:GFP transgene, possibly metaphocytes (Lin et al., 2019), were evident (e.g. Figure 7D,F,H). In *irf8* mutants, the distal segment of the transected PLLN was degraded and cleared on the same timeline as wild-type siblings (Figure 7I and J). By 1 dpt, there was significant axonal regrowth in *irf8* mutants (Figure 7F and K), and the axons had regrown fully by 2 dpt (Figure 7H and L). These results indicate that macrophages are not required for axonal regrowth after mechanical injury of the larval PLLN.

**Figure 7.**
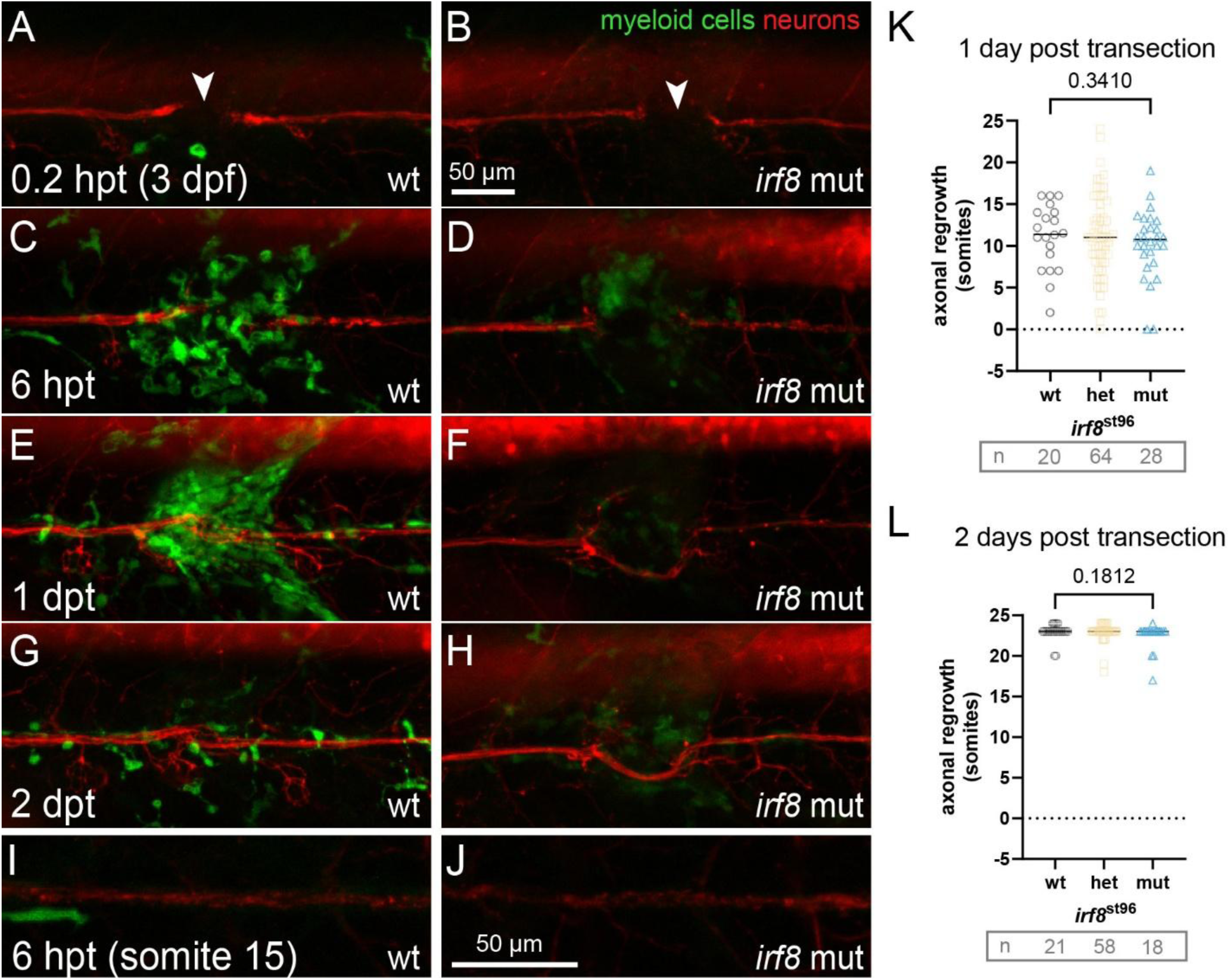
Normal axonal regrowth in *irf8* mutants lacking macrophages. (A-J). Time course of wild type and homozygous mutant *irf8* siblings expressing NBT:dsRed (neurons) and mpeg:GFP (myeloid cells). The PLLN was injured at 3 dpf, and the time points of the images are indicated. (A, C, E, G) In a wild type animal imaged at the injury site at different stages, many macrophages were apparent and the distal axons were fragmented at 6 hpt. By 2 dpt, fewer macrophages were present and the axons were regrown. (B, D, F, H). In the *irf8* mutant sibling, very few weakly GFP-expressing cells were evident. Axon fragmentation and regrowth on the same timescale as in the control sibling. (I, J) Images from the same wildtype and *irf8* mutant siblings showing axonal debris distal to the injury site. Genotypes were determined by PCR after imaging. (K, L) Quantification of axonal regrowth at 1 dpt and 2 dpt in the indicated genotypes. Lines indicate the median. Data from four independent experiments, n=animals. Mann-Whitney test used for comparisons.

## Discussion

Building on previous studies of nerve repair in zebrafish, our analysis indicates that mechanical nerve injury triggers the formation of repair Schwann cells that display many characteristics previously defined in their mammal counterparts. In adult mammals, nerve injury stimulates formation of repair Schwann cells that perform several key activities, including down-regulation of myelin genes and lengthening to form a path for regrowing axons (Jessen and Mirsky, 2016). Injured axons regrow much more rapidly in zebrafish larvae than in adult rodents, but our analysis together with prior studies indicate that many features of repair Schwann cells are nonetheless conserved between the two systems.

Mechanical transection of the nerve leads to rapid fragmentation and clearance of the distal axonal segment, which occurs in about 5 hours in the zebrafish larval PLLN (Figure 1E; Villegas et al., 2012; Ceci et al., 2014). Schwann cells begin to extend processes into the injury site by 6 hours post transection (Figure 6; Ceci et al. 2014; Xiao et al. 2015). By 1 day post-transection, distal Schwann have doubled their length (Figure 2E, F) and lost expression of the mbp:GFP reporter and Myelin Basic Protein (Figure 2D; Ceci et al., 2014). At this time, there is significant axonal regrowth across the injury site in almost all wildtype animals, and properly routed axons are growing in close association with distal repair cells (which can be distinguished by persistent changes in foxd3:GFP reporter expression; Figure 2). In wildtype larvae 3 days after transection, the axons are fully regrown and remyelination is well underway (Figures 1 and 2; Ceci et al., 2014). Thus nerve injury triggers a similar sequence of responses in larval zebrafish and adult mammals (Coleman and Freeman, 2010; Villegas et al., 2012; Gomez-Sanchez et al., 2017; Brosius Lutz et al., 2022). The combined results of studies in both systems show that the key features of repair Schwann cells are conserved.

Our analysis of the zebrafish larval system supports two key conclusions about the functions of Schwann cells at different positions within the regrowing nerve. First, proximal Schwann cells are not sufficient to direct regrowing axons across the injury site, although they can follow axons that regrow along aberrant paths (Figure 2). Using nerves in which Schwann cells populated only the proximal segment, we found that axonal regrowth failed after a distal injury (Figure 5C). There is evidence in mammals that proximal Schwann cells may lead axonal branches regrowing through the injury site from the proximal nerve stump (Cattin et al., 2015). In contrast, our analysis indicates that proximal Schwann cells often associate with misrouted axons (Figure 2A-C), which may take circuitous paths to correct targets (i.e. neuromasts). This difference may reflect the different sizes of the nerve gap, which is 15-30 times longer in adult mice than larval zebrafish (Cattin et al., 2015). In any case, our results emphasize the essential role of distal Schwann cells in guiding and stimulating axonal regrowth after mechanical injury (Figure 2A-C; Figure 5 D, E; Figure 6)(Xiao et al., 2015).

Second, a single repair Schwann cells distal to the injury site can not only direct regrowing proximal axons across the injury site but is also sufficient to allow extensive regrowth of axons through a region that is entirely devoid of Schwann cells. The key evidence supporting this conclusion derives from analysis of nerves partially populated with Schwann cells, in which the injury is positioned so that only one Schwann cell is distal to the point of transection (Figure 5 D, F, G). In these cases, proximal axons can cross the injury site and grow posteriorly over long distances that are not and have never been populated by Schwann cells. Timelapse observation also emphasizes the activity of Schwann cells immediately distal to the injury site: soon after distal Schwann cell contact, an extending proximal axon redirected toward the correct path and subsequently accelerated growth (Figure 6). Our conclusion about the necessity of distal Schwan cells contrasts with a study that addressed the role of distal Schwann cells in 1 month old zebrafish (Graciarena et al., 2014). Those authors laser-ablated cells immediately distal to the nerve cut and found that axons can regrow through the site in which distal Schwann cells had been ablated. It is possible that the laser ablation did not completely remove substrates or guidance cues that were produced by distal Schwann cells before they were ablated.

Our analyses also show that axons can regrow through regions devoid of Schwann cells after they successfully navigate the injury site (Figure 5D, F, G). This parallels the ability of the axons to extend along the correct path during their developmental outgrowth without Schwann cells (Figure 3B, C)(Gilmour et al., 2002; Lyons et al., 2005; Perlin et al., 2011). Because our model involves injury less than two days after initial nerve outgrowth, it is possible that developmental cues persist and guide the regrowing posterior lateral line nerve (PLLN) axons; further work on the guidance mechanisms is required to address this point. Our results also appear to contrast with the prevailing view from mammalian studies that repair Schwann cells and the Büngner bands they form are required along the entire length of the distal nerve segment (Jessen and Mirsky, 2016). This seeming discrepancy may reflect the different scales of the two systems—the zebrafish PLLN at 6 dpf is roughly 10 times shorter than the adult mouse sciatic nerve— and also the possible persistence of developmental cues mentioned above.

## Role of macrophages in axonal regrowth

By analyzing *irf8* mutants in our larval PLLN injury model, we found that, surprisingly, macrophages were not required for debris clearance or axonal regrowth. Prior studies in zebrafish and mammals show that macrophages phagocytose axonal debris after injury, but the requirement for macrophages is less clear. For example, one study of larval zebrafish nerve injury showed that depletion of macrophages slowed debris clearance by a few hours (Villegas et al., 2012), and another reported that epidermal cells clear sensory axonal debris with no involvement of macrophages (Rasmussen et al., 2015). Moreover, depletion of macrophages in mouse models showed only slight changes in myelin debris clearance (Perry et al., 1995), and that axonal regrowth was somewhat diminished (Dailey et al., 1998). Although we found no requirement for macrophages in regeneration of the larval PLLN, it remains important to test the possible roles of macrophages in other nerve injury contexts. For example, it has been proposed that macrophages are important to bridge large gaps in transected mammalian nerves (Cattin et al., 2015), which may not be necessary with more limited lesions.

## Limitations

Our analysis focused entirely on mechanical injury of the PLLN in early larval zebrafish. Comparisons among vertebrate models provide evidence that the functions of repair Schwann cells are conserved, but there may be important differences in role of repair Schwann cells and other cell types in different nerves, injury types, stages, or species. For example, developmental axon guidance cues may still be operative and involved in axonal regrowth at the stages we analyzed, and the path length of axonal regrowth in our injury model is significantly shorter than in mammalian models and clinical settings. Identification of the pathways controlling the formation, function, and redifferentation of repair Schwann cells will be required to determine if there are important differences in repair Schwann cells operating in different circumstances or species. In addition, we did not assess functional recovery or long-term maintenance of the lateral line system, so repair Schwann cells and other cell types may have important roles in nerve repair that were not evident from our transgenic reporter analysis.

## Supplemental movie 1

In a wild type animal with labeled neurons (NBT:dsRed, red) and Schwann cells (foxD3:GFP, green), the PLLN was transected as described above at 3 dpf. This timelapse of the injury site begins 8 hours post-transection and runs for 25 hours, with an interval of 10 minutes between frames. The distal Schwann cell at the injury site makes contact with a proximal axon projecting ventrally at 5.3 hours into the movie (frame 32). After contact the axon proceeded along the distal Schwann cell process until it reached the main distal nerve segment, and regained the correct course. The axon analyzed has projected down the main trunk of the PLLN and passed out the field of view at 15 hours (frame 92). Single axons are thin and dim in comparison to the intact nerve; fine processes that are not evident in the timelapse as shown can be resolved after brightness is adjusted (e.g. Figure 6C insert).

## Acknowledgements

This work was supported by the National Institutes of Health/NINDS (R35 NS111584 to W.S.T.). We thank members of our lab for helpful discussion and Tuky Reyes and Chenelle Hill for fish care. W.S.T. is the Mary and Dr. Salim Shelby Professor at Stanford University.

---

**Figure S1.**
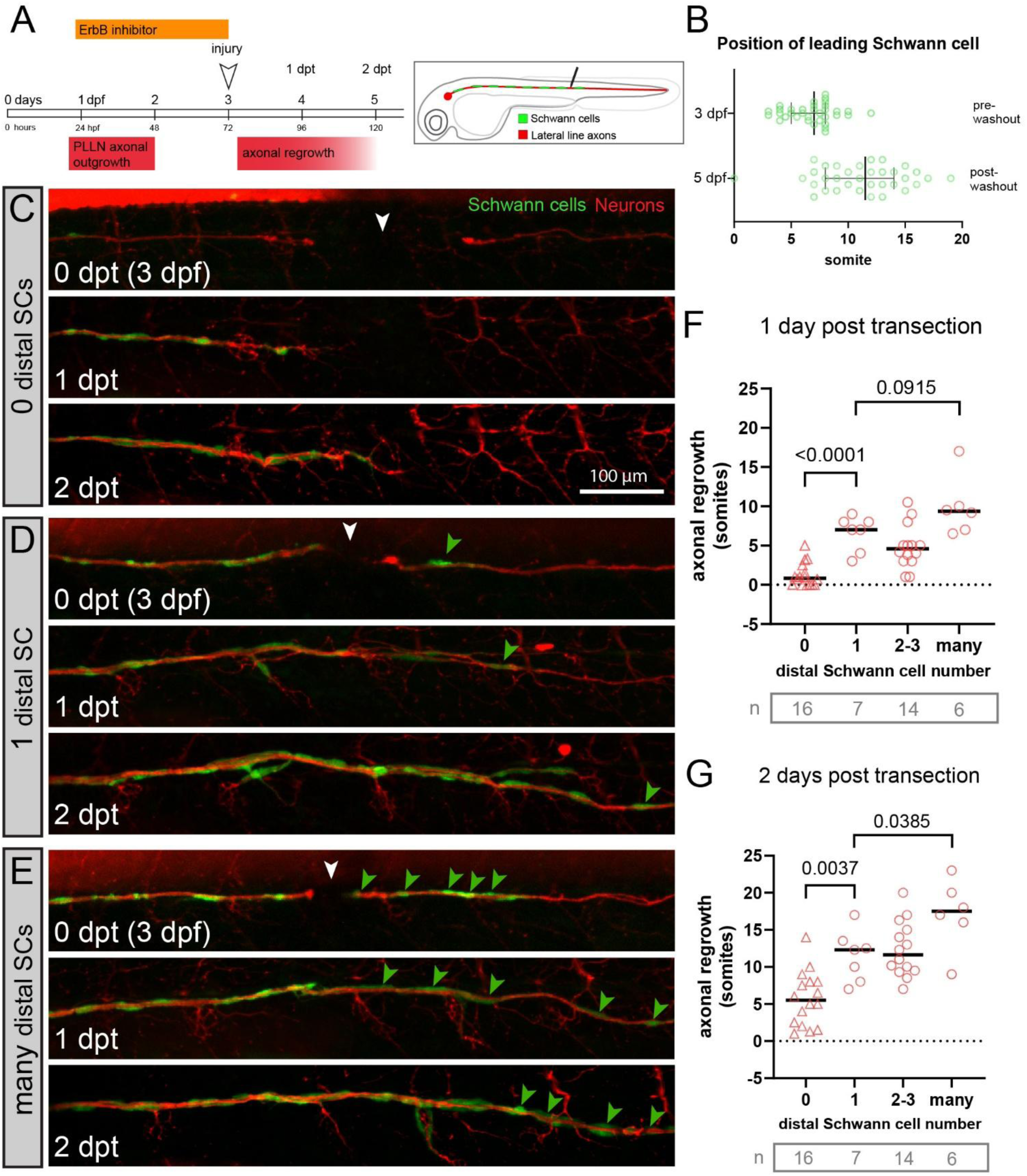
After ErbB inhibitor washout, a single Schwann cell was sufficient for significant axonal regrowth after nerve transection. (A) Timeline of ErbB inhibitor treatment to generate nerves partially covered in Schwann cells. ErbB inhibitor is removed post-injury. (A’) Transection paradigm to generate an injury site with a single distal Schwann cell. (B) While Schwann cell migration was disrupted by ErbB inhibition, after injury and ErbB inhibitor removal, Schwann cells migrated an average of ∼5 somites further posterior. (C-E) Animals expressing foxd3:GFP (Schwann cells, green) and NBT:dsRed (axons, red) were imaged at different time points from injury through 2 days of regrowth (white arrowhead at injury site, green arrowheads mark distal Schwann cells) (C) Without a distal Schwann cell, axons stalled at the injury site at 1 dpt. Only modest growth was observed at 2 dpt. (D) With a single distal Schwann cell, significant axonal growth was apparent at 1 dpt and continued at 2 dpt. (E) Without ErbB inhibitor and with many distal Schwann cells, axonal regrowth was slightly more than when when one distal Schwann cell was present. Schwann cell migration resumed after ErbB inhibitor was removed, as illustrated by the more posterior positions of indicated distal Schwann cells. (F,G) Quantification of axonal regrowth at 1 dpt and 2 dpt. Data from three independent experiments, n=animals. Lines indicate the median. Mann-Whitney test used for comparisons.

